# Treasure: A Sensitive Pipeline for Species-Level and Functional Microbiome Profiling

**DOI:** 10.1101/2025.09.03.674071

**Authors:** Daniel de Souza Avelar, Eliel Barbosa Teixeira, Samir Mansour Moraes Casseb, Fabiano Cordeiro Moreira, Paulo Pimentel de Assumpção

## Abstract

Next Generation Sequencing (NGS) methods, such as 16S rRNA amplicon sequencing and Whole Genome Sequencing (WGS), enable taxonomic analyses but have limitations. This project proposes the development of a computational tool capable of performing functional analysis of the most abundant microorganisms within a microbiome based on taxonomic analysis. The proposed method integrates the tools Kraken, Gffread, and Salmon. Compared to Samsa 2, a commonly used pipeline for RNA-Seq Total samples, the new approach demonstrated superior performance across all evaluated scenarios (p < 0.01). The tool aims to functionally characterize the microbiome of regions affected by Gastric Cancer (GC) and adjacent areas, assess associations between survival, expression/abundance, and identify potential microbial biomarkers for GC.

## 1. INTRODUCTION

In recent years, the advent of Next-Generation Sequencing (NGS) has allowed new perspectives for studies in the field of microbiology (1) based on meta-omics analyses (metagenomics, metatranscriptomics, metaproteomics and metabolomics), which are approaches that aim to understand the human microbiomes in diseases (2, 3, 4), since they help in the characterization of microorganisms, strains identification, expression patterns and in the elucidation of metabolic pathways (5).

The method usually used in these analyses is sequencing hypervariable regions (v1-v9) of the 16S rRNA gene for taxonomy and phylogeny studies (6). However, the most accurate classification given by this method is frequently at the genus level, and it is not ideal for characterizing the functional profile of the investigated community since all functions are inferred by biological pathways previously associated with these bacteria. (7). Furthermore, as 16S rRNA is only in bacteria and archaea, it disregards other microorganisms present in the microbiome, such as fungi and viruses, that may contribute to the microbiome dynamics (8). On the other hand, metagenomics allows identifying bacteria down to the species level and investigating all microorganisms.

Metatranscriptomics is the overall analysis of the sample transcripts (9), a knowledge that allows inferences about microbial activity through observations of gene expression patterns (10) and, in turn, contributes to clarifying the paradigm: Presence vs. Activity (9, 11). Therefore, it points to those transcriptionally active in a given spatial and temporal context in host-pathogen interactions (12). Furthermore, for clinical microbiology, identifying functionally active bacteria is essential to indicate disease-causing organisms (13,14).

Based on the scenario mentioned earlier, using protocols and tools to guarantee the quality and reliability of the data generated is essential. However, since these samples are sequenced with unknown and dynamic scenarios, it is common to find several pipelines related to the metatranscriptome. Therefore, information such as depth, coverage, and tools for analyzing the sequencing generated data are diverse (15). Currently, the following pipelines are available: MetaTrans (16), COMAN (17), FMAP (18), SAMSA2 (19), HUMANN2 (20), SqueezeMeta (21), IMP (22) and MOSCA (23), used for RNA-seq data analysis.

SAMSA2 is the most used pipeline due to its fast speed since it uses DIAMOND internally (19). The DIAMOND aligner is an open-source program that compares sequencing readings against a protein database (24), which is why it is integrated into most metagenomic data analysis protocols, mainly due to its processing speed compared to existing ones.

However, DIAMOND’s database can lead to errors during the annotation phase since converting nucleotide sequences into amino acids can generate multi-mapping. The multi-mapping allows the alignment of the readings in more than one genomic location (25). Thus, it becomes, in fact, something harmful when considering the redundancy of the genetic code; that is, the conversion into amino acids can generate changes in the sequence nucleotide mutation, which can result in missense, nonsense or silent mutations.

Although these modifications are silent, resulting in the same amino acid (26), it becomes a problem in metagenomic analysis because the alignment from amino acids will have a similar identity, even in bacteria with different nucleotide sequences.

In contrast to the presented problem, the present study introduces the Treasure tool alternatively. Treasure uses Salmon (27) and Kraken 2 (28) as sequence aligners, the former responsible for expression analysis at the nucleotide level; the second to produce taxonomic classification. Therefore, the merge of Salmon and Kraken 2 in Treasure is a differential in the metatranscriptomics data analysis.

## 2. MATERIALS AND METHODS

### 2.1 SOFTWARE DEVELOPMENT

Like other tools that use next-generation sequencing data, Treasure is grouped modularly. These modules are named as follows: Configuration, alignment, meta-alignment and update.

#### 2.1.1 CONFIGURATION

The configuration module, known as “config”, establishes parameters for the pipeline execution. During configuration, user inputs:

- Sample location
- Project name (determines output file prefix)
- Output folder name
- Number of threads
- Sample type: Single-End (SE) or Paired-End (PE)

For PE samples, users specify processing preferences: paired and unpaired or only paired. Unpaired reads are stored in separated files. The program also supports compressed.fastq.gz samples. After proper error checking, the program finishes the mode and saves the information in a configuration file in JSON format. The configuration file uses “.config.dio” as its extension.

#### 2.1.2 ALIGNMENT

The alignment module (“align”) uses *Kraken 2* (28) to align and detect the microorganisms present in the sample. The *Kraken 2* database comprises the human genome, bacteria, viruses and archaea. The user must download the database since it is a mandatory program resource (https://benlangmead.github.io/aws-indexes/k2). After the execution of each sample, the result of all is standardized and attached in a single file (.taxReport.dio).

#### 2.1.3 META-ALIGNMENT

During the meta-alignment stage (“metaAlign”), the genomes of the most abundant microorganisms will be acquired, and the reads will undergo realignment using Salmon to obtain the gene expression profiles for each organism. Users have the option to specify the analysis target (Bacteria or viruses) and choose the taxonomic level (species or genus). The number of predominant microorganisms analyzed can be adjusted by the user, with a default setting of 10.

In a hypothetical scenario, a metatranscriptomic experiment is conducted on a total of 40 samples, structured within a case–control design. The experimental groups are defined as follows:

- Case: S1-S20;
- Control: S21-S40.

The computation of the most prevalent microorganisms can be achieved through three sub-modes: Unified, Metadata, and One-to-one. Utilizing these sub-modes allows us to answer specific questions within this experimental framework.

- Unified: This is the default method of the program, in the absence of any argument, involving using all samples as a reference. In the hypothetical experiment, all 40 samples would be utilized to calculate the N predominant microorganisms.
- Metadata: When the user provides a metadata file, delineating the samples into groups, such as in a case-control scenario, the set of N most abundant microorganisms is extracted based on the specified group. In the hypothetical experiment, if the “case” group were selected, the predominant microorganisms would be calculated using this set of samples as a reference. It’s important to note that even if the “case” group is selected, all samples will still be aligned using this reference set.
- One-to-one: Computes the most abundant microorganisms for each sample individually, adopting a distinct alignment strategy where each sample is aligned with its respective reference genome containing the most abundant microorganisms. In this approach, samples are not categorized into groups like case or control; rather, microorganism abundance is assessed on a per-sample basis. To prevent redundant downloads, the program identifies intersections between microorganisms within samples. Subsequently, a composite transcriptome is generated for each sample, facilitating alignment with its respective transcriptome.

The program calculates the most abundant microorganisms using the following algorithm:

1. *For each taxon:*
  a. *Sum the reads of this taxon in all analyzed samples;*
2. *Rearrange the values of these sums in a descending order*.

The first N values are the most abundant microorganisms. The same reasoning holds for an alignment at the genus level.

Following the identification of the most abundant species, the program retrieves the links facilitating access to their genomes from a refined version of the RefSeq database for bacteria and viruses. This database contains both genomic sequences (.fna) and genomic annotation files (.gff). If a species is not found during the initial download, the program offers the user the option to update the database with the latest RefSeq data. If the species is absent from both databases, the user can input the taxonomic id of a species that more accurately represents the microorganism under analysis. Alternatively, if preferred, the program can proceed without including that species.

Upon acquiring the genomes, references, and annotations, the Gffread tool (31) is utilized to extract the transcripts FASTA file for each species. Subsequently, these files are consolidated to form a unified genome, stored within the project folder for potential future reference or independent alignment purposes. Utilizing the obtained reference transcriptome, the Salmon aligner (27) is employed to perform gene quantification. Salmon was selected for its capability to discern isoforms and offer precise nucleotide-level alignment. Additionally, Salmon can run with Single-End (SE) and Paired-End (PE) samples. If the user has opted to work with unpaired reads during the configuration phase, these will be aligned separately as an SE sample using the file generated in the preceding step.

##### 2.1.3.1 INTRAGENUS REPRESENTATION SCORE

For analysis at the genus level, the selection of representative species is required. Therefore, the Intragenus Representation Score (IRS) was devised, which for each species within a genus is calculated by:

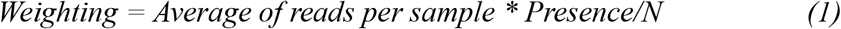

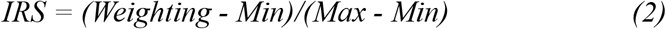

The initial step in this calculation is the Weighting [1], which is determined by the species’ mean presence per sample. Subsequently, it undergoes normalization between 0 and 1 using the described min-max normalization method. Species that achieve the threshold of 0.7 (default setting) for the IRS will serve as references for the assessed genus. This threshold is adjustable by the user.

In this manner, for analysis at the genus level, all reads will be considered, and the species that best represent the genus (based on the IRS) will have their genomes downloaded.

In Table 1, depicting four fictitious species, we assess which one(s) best represent(s) the genus. While considering the highest value in a single sample suggests choosing species sp1, its absence in samples 1 and 2 may indicate potential contamination. Although sp4 could be chosen by summing all reads in the sample set, its absence in sample 3 and exaggerated value in sample 2 might introduce bias towards the mean. While sp3 has a higher weighting value than any other species, it remains smaller than T2. However, instead of selecting only species T2, the IRS metric, associated with a threshold of 0.7, permits the selection of multiple species to represent the evaluated genus. Therefore, both sp2 and sp3 should be elected as representatives of this hypothetical genus.

**Table 1.** Weighting calculation to choose the microorganism that best represents a hypothetical genus.

| Sample | S1 | S2 | S3 | Mean | Pres/N | Weighting | Score |
| --- | --- | --- | --- | --- | --- | --- | --- |
| Espécie |  |  |  |  |  |  |  |
| sp1 | 0 | 0 | 20 | 6,66 | 1/3 | 2,22 | 0 |
| **sp2** | 10 | 2 | 7 | 6,33 | 3/3 | 6,33 | **1** |
| **sp3** | 6 | 6 | 6 | 6 | 3/3 | 6 | **0,91** |
| sp4 | 5 | 17 | 0 | 7,33 | 2/3 | 4,88 | 0,64 |

#### 2.1.4 UPDATE

The update mode (“updateDB”) downloads the latest RefSeq database, which then coexists alongside the original database of the program. When this option is activated during the download of genomes for species identified during the analysis, Treasure queries both the original and updated databases. Users have the option to request the update independently or during the download of species that are absent from the original database of the program.

### 2.2 VALIDATION

#### 2.2.1 COMPARISON AGAINST SAMSA 2

To benchmark against Samsa 2, four mock community types (water, soil, feces, and tissue) were constructed to assess tool performance across distinct scenarios. For each sample type, a literature survey was conducted to identify the most prevalent taxa within each domain (Supplementary Material, Appendix I).

Representative genomes were retrieved from the RefSeq database and combined to generate composite genomes, one per sample type. Detailed taxonomic compositions are provided in Supplementary Material, Appendix II.

From each composite genome, datasets of 10 million reads were simulated using the “randomReads.sh” script from BBTools (32). The resulting community compositions are reported by taxonomic group in the Results section. For every read, the source species, a unique gene identifier, and, when available, the shared gene symbol were annotated in the read header for downstream analyses.

Tool performance was quantified using a confusion matrix, with true positives (TP), true negatives (TN), false positives (FP), and false negatives (FN) defined per class. Standard performance metrics were then computed for each category.

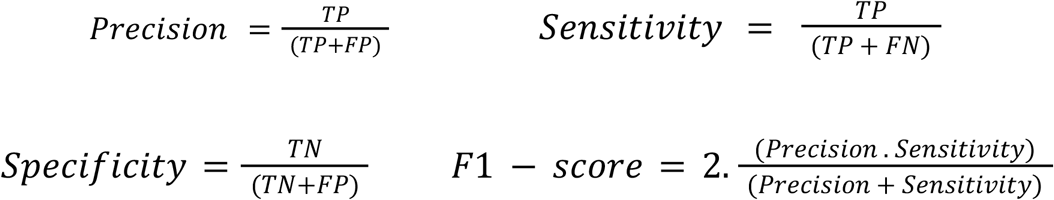

Tool performance was evaluated across three levels: species, taxonomic group, and gene. For species detection, each species present in the mock community was represented as a row/class in the confusion matrix, with columns corresponding to species identified by the tools. Species not detected by either tool were omitted from the plot, with the number of omissions reported in the Results section.

For functional analysis, only genes with shared symbols were included, while unique genes were excluded due to variability across microorganisms and genomic annotations. An illustrative case is provided in Table 2.

**Table 2.**
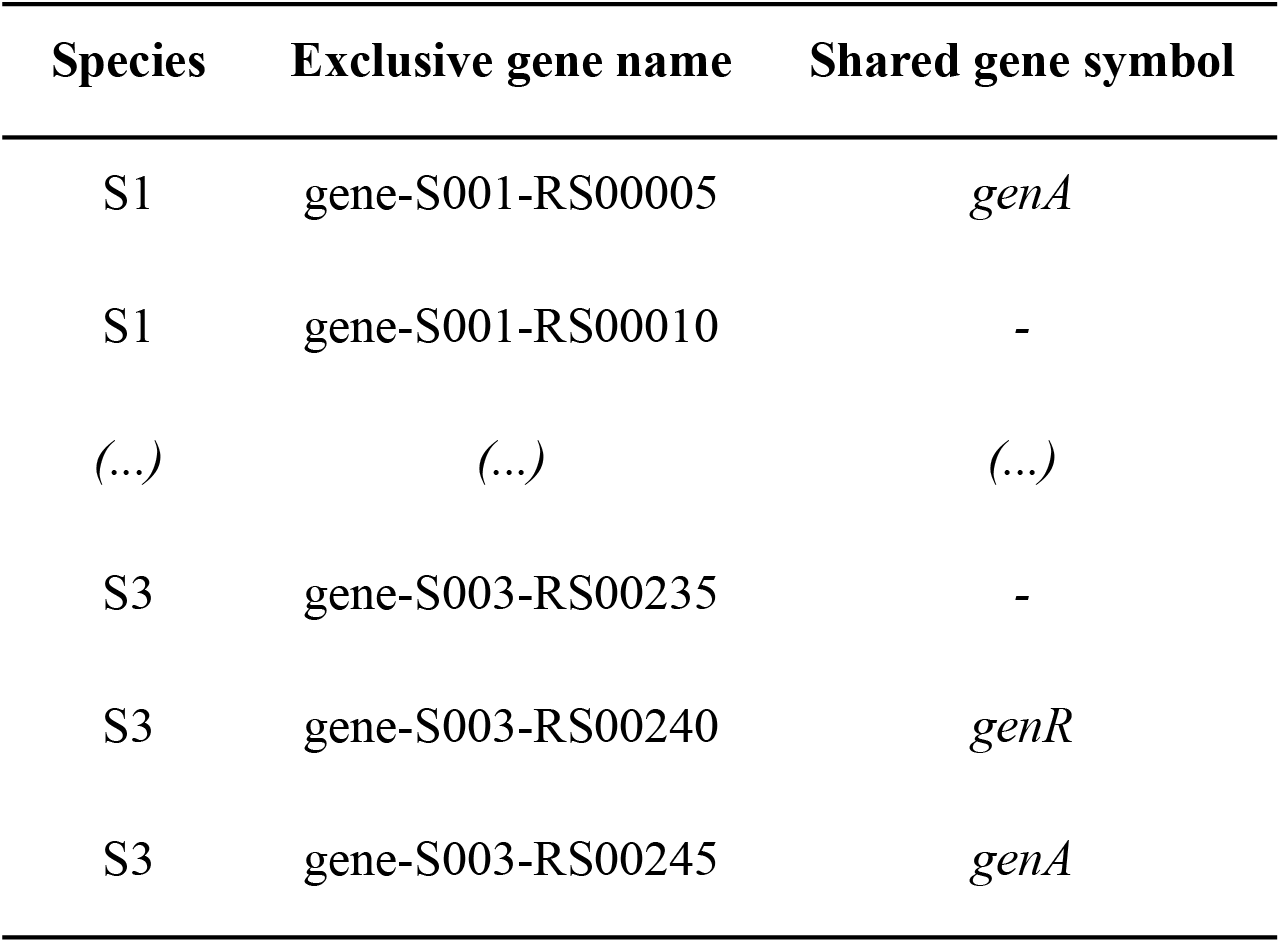
Example of shared genes and their evaluation by the tools.

| Species | Exclusive gene name | Shared gene symbol |
| --- | --- | --- |
| S1 | gene-S001-RS00005 | *genA* |
| S1 | gene-S001-RS00010 | - |
| (...) | (...) | (...) |
| S3 | gene-S003-RS00235 | - |
| S3 | gene-S003-RS00240 | *genR* |
| S3 | gene-S003-RS00245 | *genA* |

Given that exclusive gene annotations do not necessarily follow a uniform standard across genomes of the same species or genus, shared genes represent a more reliable categorical variable for downstream analyses. In the scenario illustrated in Table 2, columns lacking shared genes were excluded, retaining only rows corresponding to genA and genR.

In addition to filtering based on shared genes, a second filter was applied, in which only genes identified by at least one tool were included in the final boxplot. The total numbers of reads and genes excluded are detailed in the text.

Each row of the confusion matrix corresponds to genes present in the simulated community, while columns represent genes identified by each tool. For the evaluation of taxonomic groups — including bacteria, viruses, archaea, fungi, and protozoa — the origin of each simulated read is known, allowing comparison between the predicted and observed taxonomic group for each tool.

The F1-score, as the harmonic mean of precision and recall, was chosen as the metric for the boxplots across all three classes. Correct identification was calculated for each class across the three described scenarios. Statistical significance between tools was assessed using a two-sided Mann–Whitney–Wilcoxon test with Bonferroni correction for multiple comparisons, considering *p* < 0.05 as the threshold for significance.

#### 2.2.2 COMPARISON WITH LITERATURE

For comparison with the literature, data from the article by Hadzega et al. were used. These breast cancer tumor and normal samples represent reads that were not aligned with the human genome (HG38). During the study, the Kraken 2 tool also was used. This data is available under accession id PRJNA751534.

In R v4.2, for the taxonomic analysis, the 30 most abundant microorganisms were calculated by each group. For the following differential analyses, the DESeq2 package was used. The taxon was considered statistically significant when p-value < 0.05 and log2FC > 1. For the differentially expressed genes analysis, the gene was considered statistically significant when p-value < 0.05 and log2FC > 2.

### 2.3 DATA AVAILABILITY

Treasure is available in GitLab through the following link: gitlab.com/avelardaniel/Treasure.

## 3 RESULTS

### 3.1 COMPARISON AGAINST SAMSA 2

Table 3 details the composition of each simulated community in terms of taxonomic groups prior to the application of the scenario-specific filters described in the Methods.

**Table 3.** Composition of mock communities.

| Taxonomic group |  | Water | Feces | Soil | Tissue |
| --- | --- | --- | --- | --- | --- |
| Bacteria | n= | 46 | 44 | 7 | 40 |
|  | % | 87,16% | 81,39% | 25,01% | 39,64% |
| Viruses | n= | 6 | 5 | 9 | 34 |
|  | % | 0,15% | 0,03% | 1,78% | 0,44% |
| Archaea | n= | 2 | 1 | 1 | 6 |
|  | % | 1,54% | 1,5% | 0,96% | 5,69% |
| Fungi | n= | 1 | - | 7 | 15 |
|  | % | 2,79% | - | 72,23% | 54,21% |
| Protozoa | n= | 1 | 4 | - | - |
|  | % | 8,33% | 17,06% | - | - |

In Figure 3, the tool developed in this study demonstrates superior performance in taxonomic identification across all tested samples. For the Water, Feces, Soil, and Tissue samples, after applying a filter that retained only taxa detected by at least one of the tools, approximately 75%, 68%, 60%, and 74% of the analyzed microorganisms were preserved, respectively.

**Figure 1.**
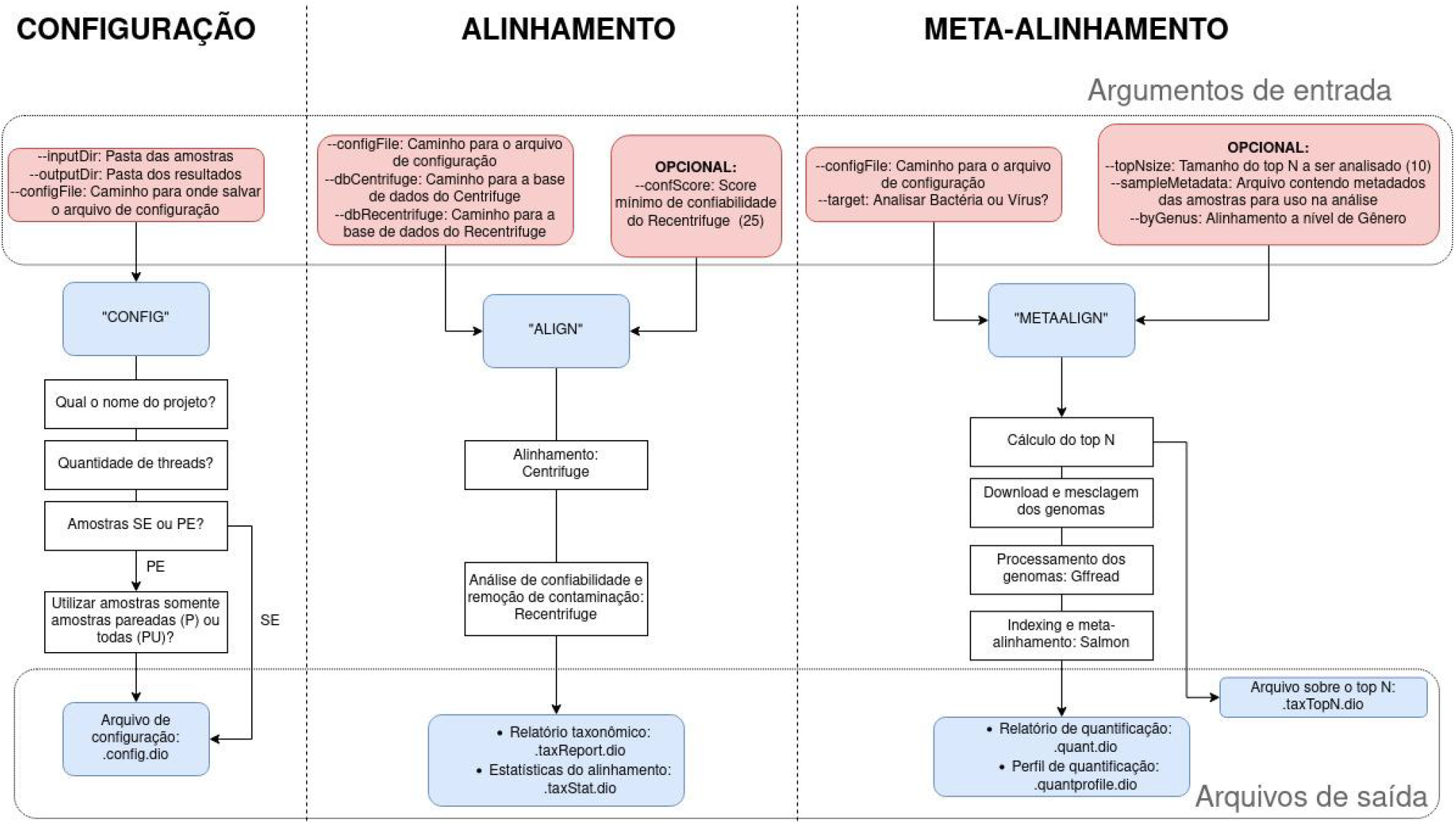
Main modules of Treasure

**Figure 2.**
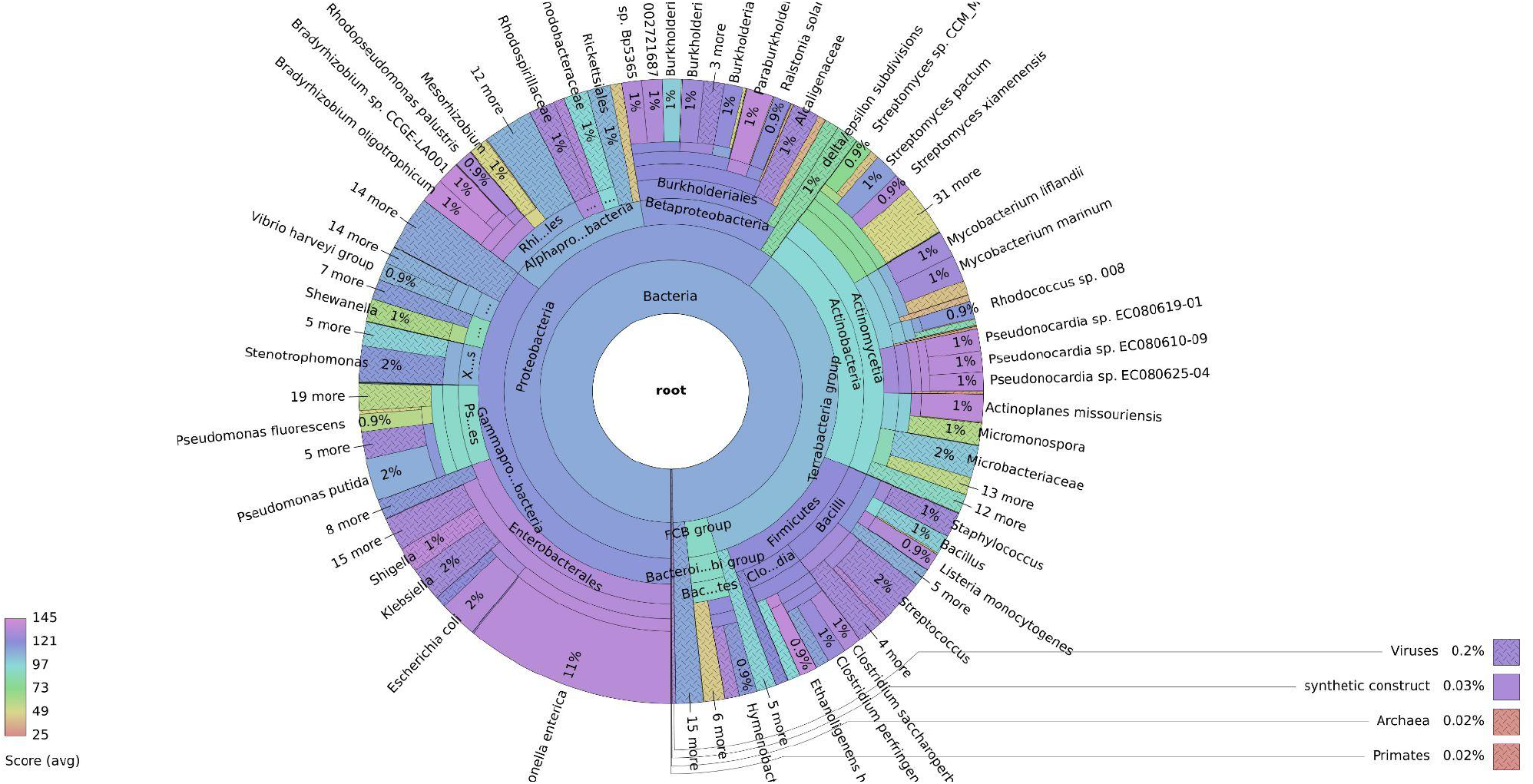
Taxonomic result generated by this tool for the mock community

**Figure 3.**
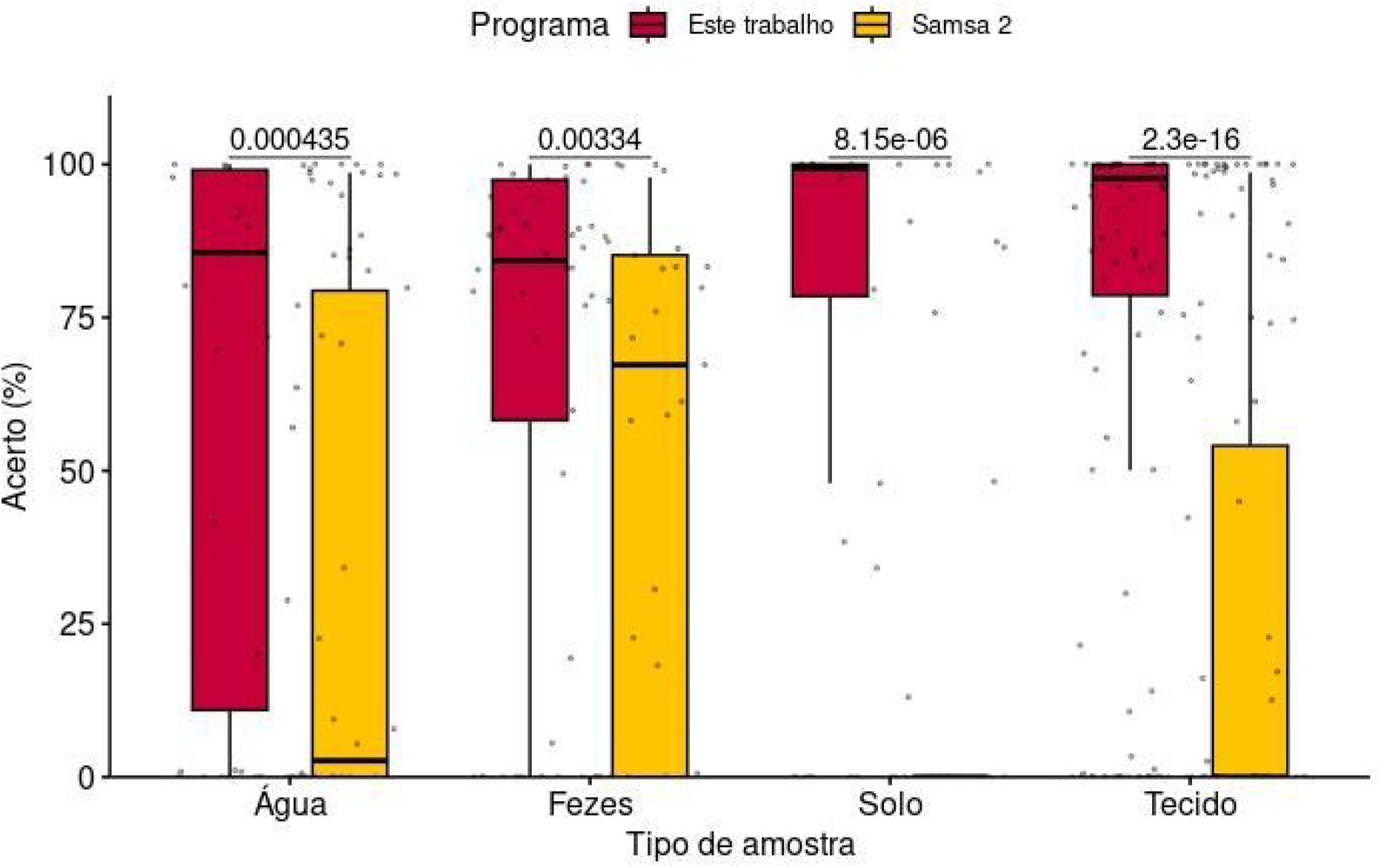
Accuracy percentage by taxon comparing the performance of the tool developed in this study with Samsa 2

The comparison shown in Figure 4 indicates that the tool is able to identify all taxonomic groups with a satisfactory percentage, except for protozoans. Samsa’s performance is similar to that of the tool for bacteria; however, it fails to detect any of the other domains. The highest performance of the tool is observed for the viral taxonomic group, followed by the bacterial group.

**Figure 4.**
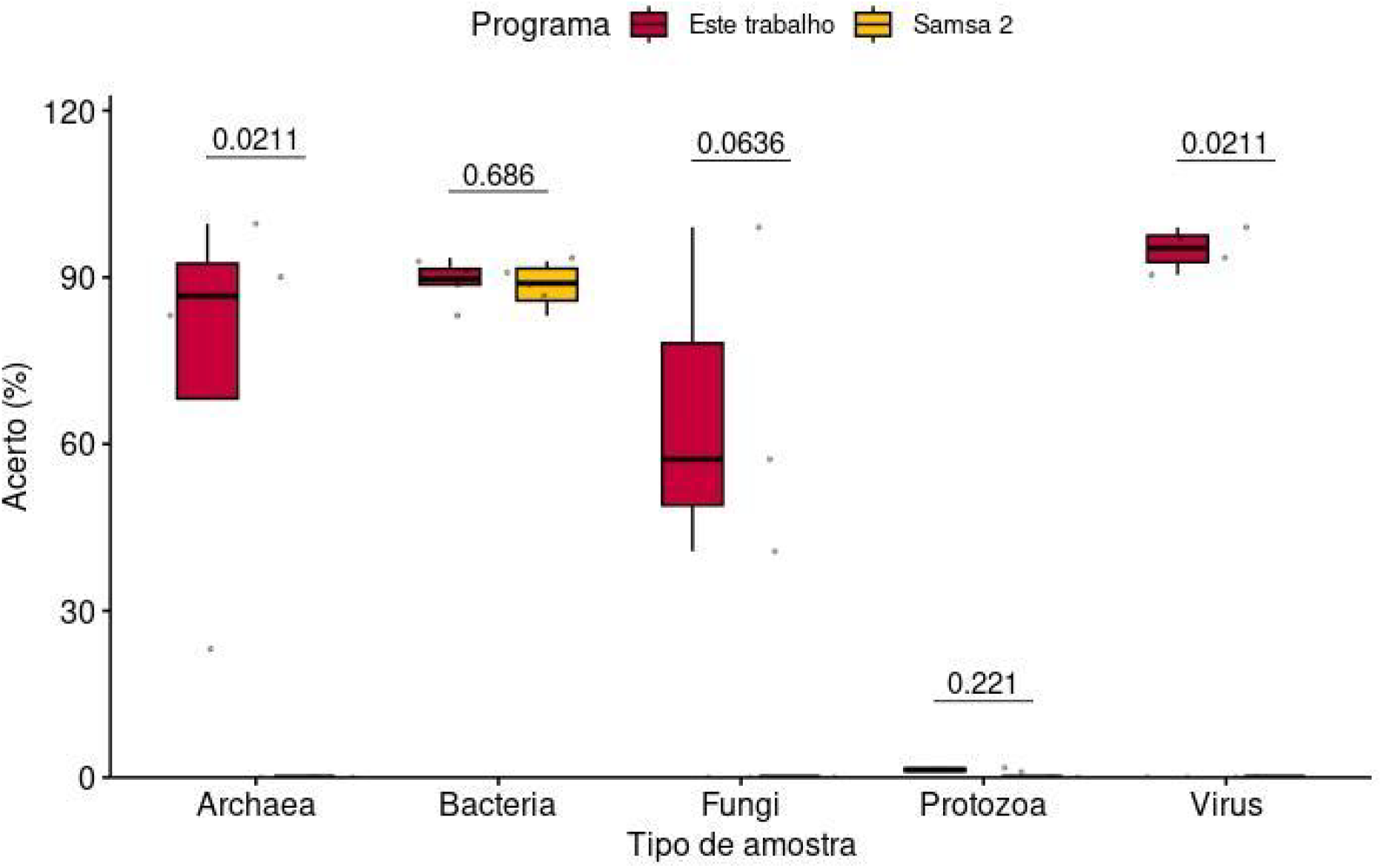
Accuracy percentage by taxonomic group

Gene-level analysis was conducted using only genes with shared symbols, excluding unique genes due to possible variations among microorganisms and genomic annotations. As noted in the methodology, not all reads from the simulated community correspond to a gene with a shared symbol; therefore, not all reads could be used in this evaluation. Additionally, as described in the methodology, only genes detected by at least one of the tools were included in this comparison. Considering the filters described above, Table 4 presents the number of genes evaluated per sample, which was used to generate the data represented in Figure 4.

**Table 4.** Gene filtering for functional-level comparison between the tools.

| **Samples** | **Reads with gene symbol** | **Total shared genes found** | **Evaluated genes** |
| --- | --- | --- | --- |
| **Tissue** | 1.246.809 | 4880 | 1313 |
| **Feces** | 2.617.677 | 5009 | 226 |
| **Water** | 2.121.133 | 4242 | 524 |
| **Soil** | 921.870 | 4842 | 1697 |

After applying the filters, an average of 17.26% of the samples remained for functional analysis. To avoid any presence-related bias, only genes detected by at least one of the tools were analyzed. Gene-level comparison, shown in Figure 5, indicates that the tool outperforms Samsa across all scenarios.

**Figure 5.**
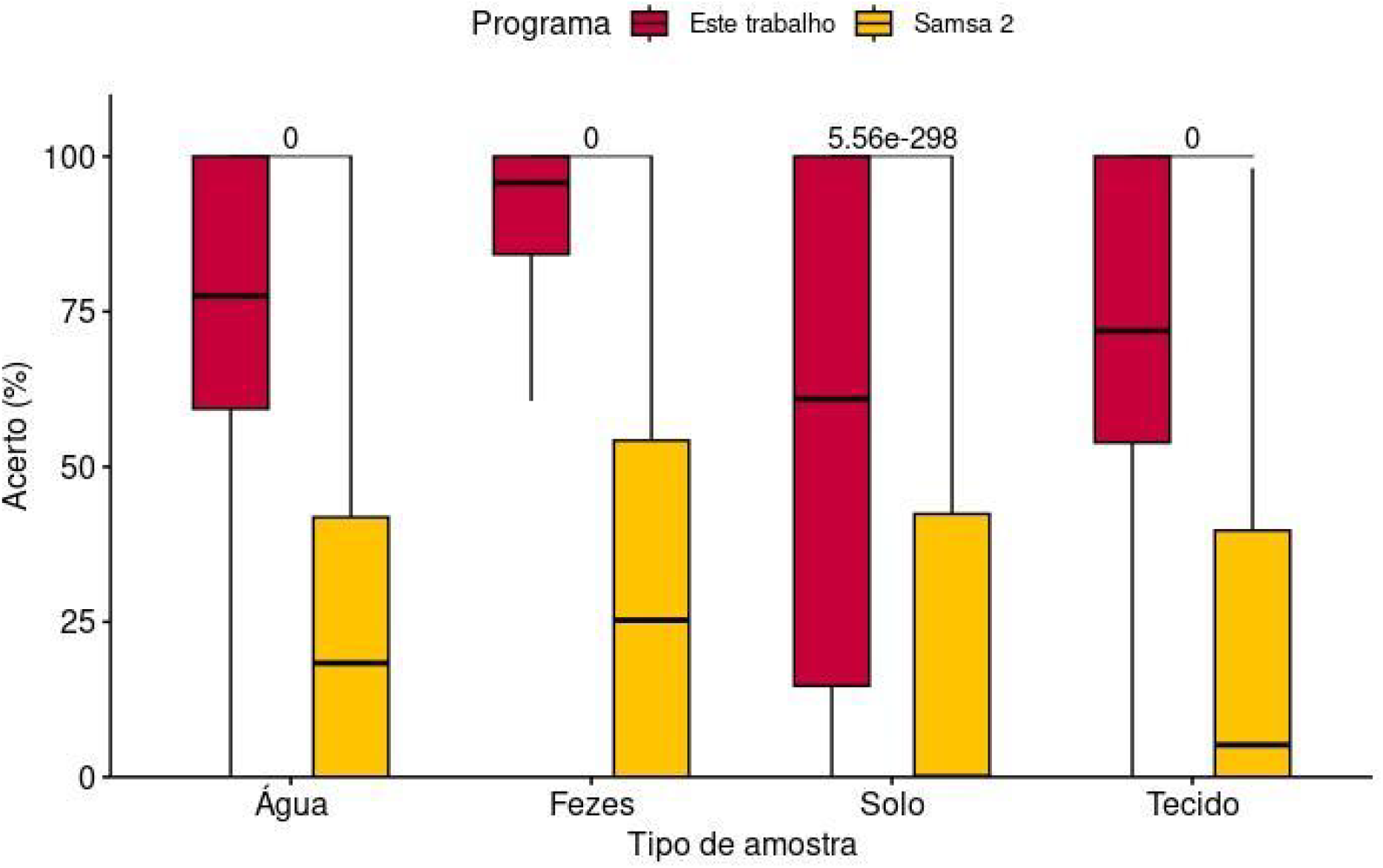
Gene-level comparison

### 3.2 COMPARISON WITH LITERATURE

For comparison with real breast tissue samples, as presented in Figures 6 and 7, the Cancer group exhibits the following taxa among the ten most abundant families, listed in descending order of abundance: *Bacilaceae, Burkholderiaceae, Clostridiaceae, Comamonadaceae, Enterobacteriaceae, Halomonodaceae, Lactobacilaceae, Microbacteriaceae, Micrococcaceae* and *Moraxellaceae*. For the Normal group we have the following order: *Bacilaceae, Bacteroidaceae, Burkholderiaceae, Clostridiaceae, Comamonadaceae, Cytophagaceae, Enterobacteriaceae, Erwiniaceae, Halomonadaceae* and *Hymemobacteriaceae*.

**Figure 6.**
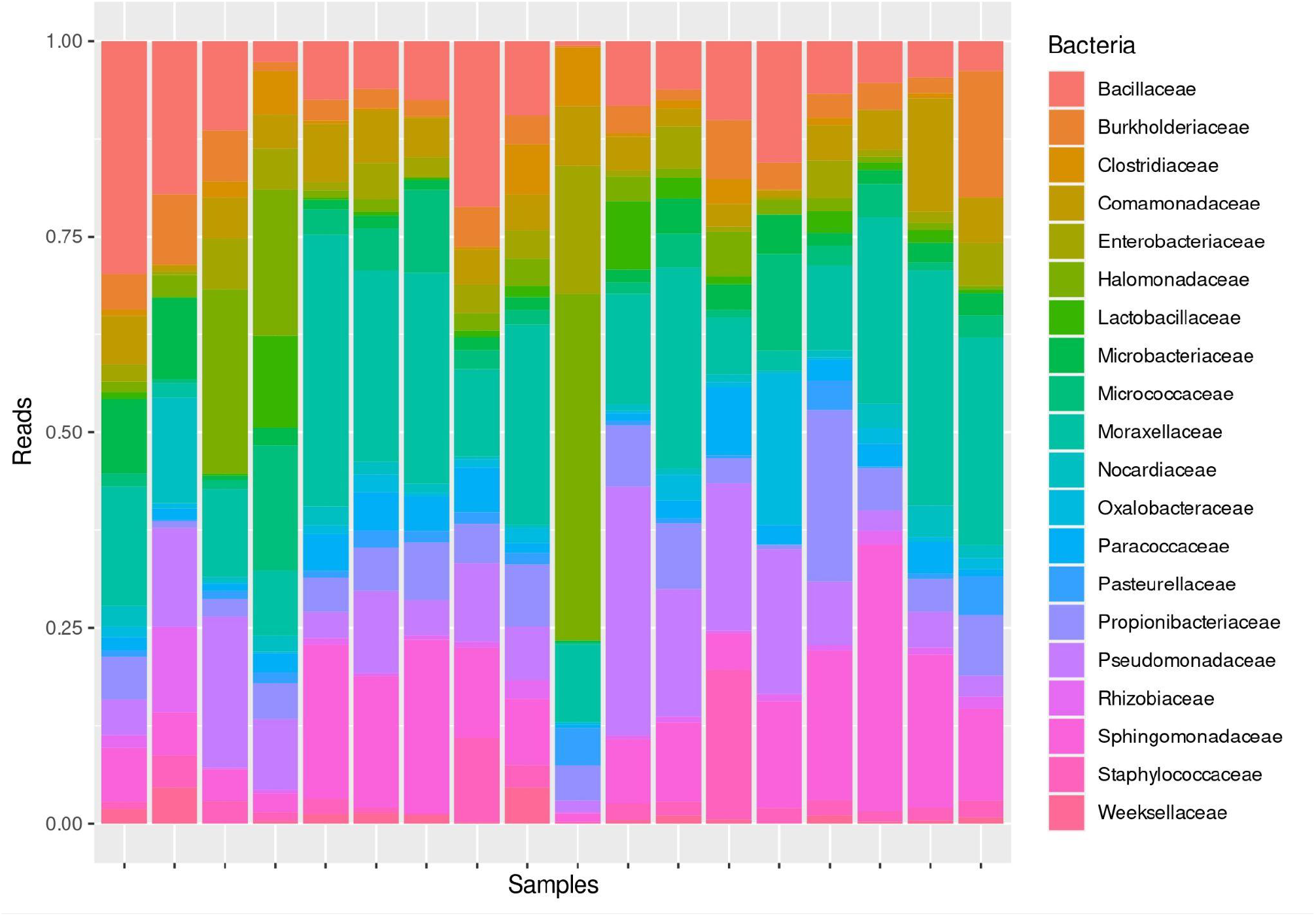
Most abundant taxonomic families in breast cancer samples

**Figure 7.**
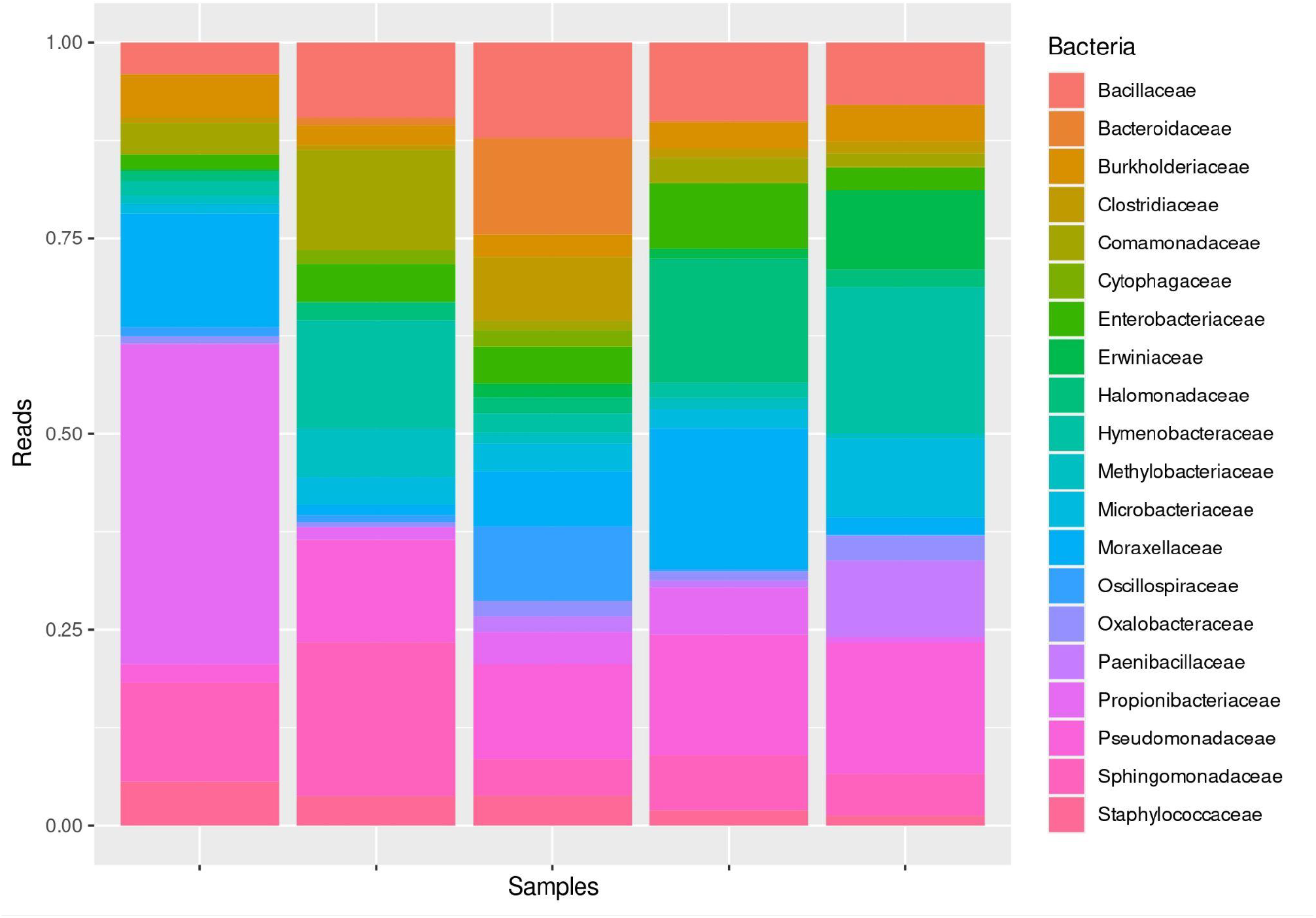
Most abundant taxonomic families in normal samples

When evaluating the differentially abundant genera by histological type, it is possible to observe, in Figure 8, a total of 22 DA families, 3 of them in the tissue affected by breast cancer and 19 in the healthy tissue.

**Figure 8.**
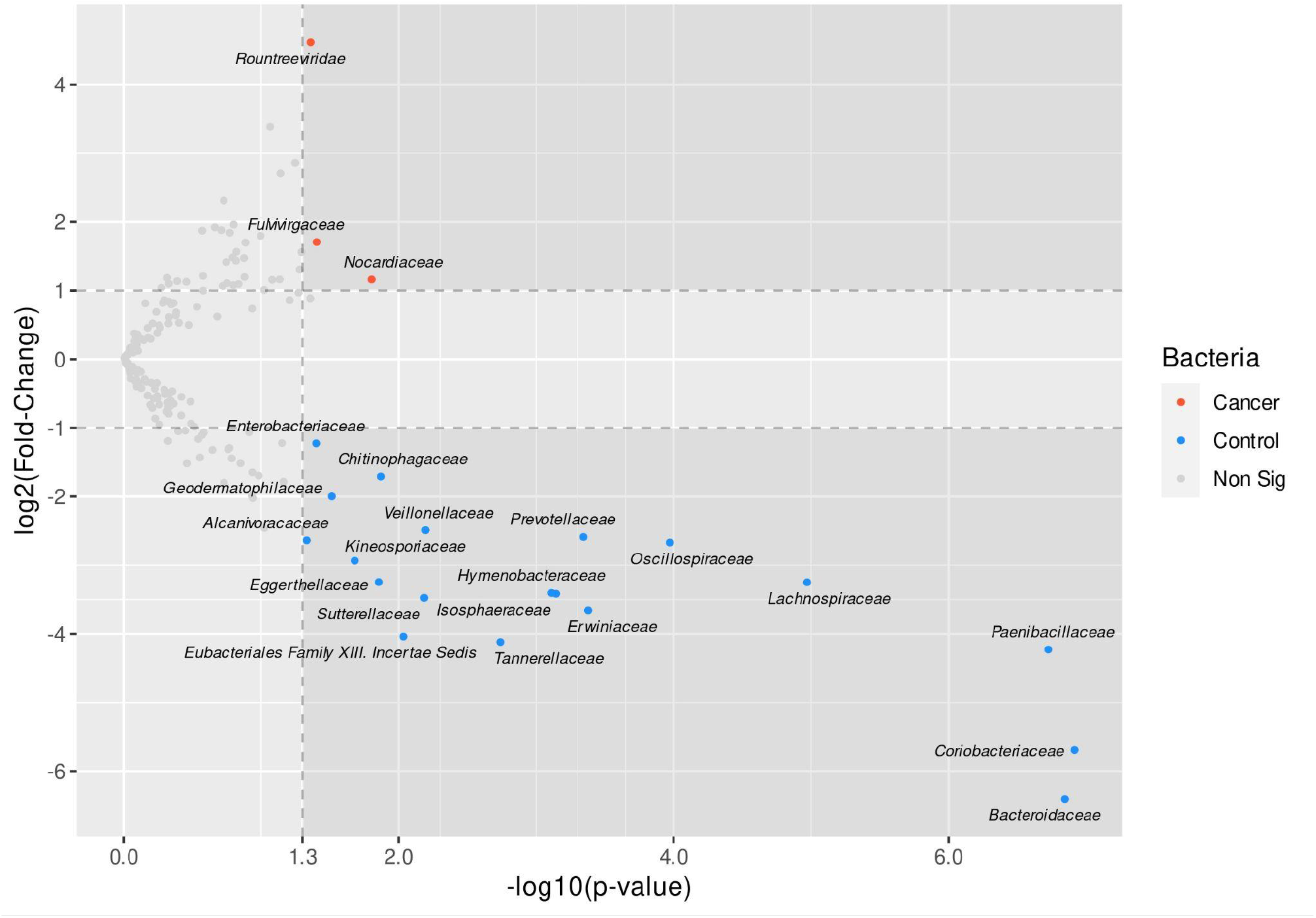
Comparison of differentially abundant genera between Cancer and Normal

By analyzing differentially expressed bacterial genes, using data from the last step of the pipeline, we found 40 DE genes, where 26 of these are found in tumor and 14 of these are found in normal tissue.

## 3. DISCUSSION

Treasure is a pipeline for integrated analysis in RNA-Seq data. Therefore, it is postulated as a potential alternative to Samsa 2 since it detects a more significant number of microorganisms’ reads than its counterpart. The tool aligns a set of samples, detects the most abundant microorganisms, and does a quantification step to calculate the genic expression of these bacteria or viruses.

The *Kraken* uses a model based on k-mers and the whole genome, making it more accurate by evaluating the genome sequence instead of specific biomarkers. In addition to using less RAM and disk space, *Kraken* still maintains the taxonomic identification benchmark when compared in the literature with other tools with a similar objective, as shown in the article by Kim et al.

Aligners with divergent methodologies, such as those operating at the nucleotide versus amino acid levels, fall under the paradigm of sensitivity versus specificity. Although amino acid-level aligners have the specificity to identify true alignments, sensitivity is reduced due to the redundancy of the genetic code, since the same amino acid can be encoded by multiple codons. Diamond, used internally in Samsa 2, is an amino acid-level aligner, whereas Kraken operates at the nucleotide level. Our tool detects a greater number of species and demonstrates superior sensitivity at the species, functional, and taxonomic levels, which explains its higher overall performance, as shown in the results and in Table 1 of the supplemental material.

The article by Hadzega et al. was used to compare with a real dataset. This work uses a methodology similar to the one proposed in this article, since it characterizes the microbiota in primary breast cancer with Total RNA-Seq samples. When analyzing data from the article by Hadzega et al. with Treasure, we observe that the results are very similar at family level, as shown in Figure 6.

A key aspect of this study is that total RNA sequencing is a valid methodology for taxonomic and functional characterization compared to 16S. We observed that the method offers notable sensitivity at the family level and enables functional characterization of the microbiota.

Given that the article uses Total RNA-Seq samples, a possible step is the functional and differential expression analysis. This brings us to Salmon, which can provide the result presented in Figure 9. Salmon is an aligner capable of quantifying gene expression based on a reference.

**Figure 9.**
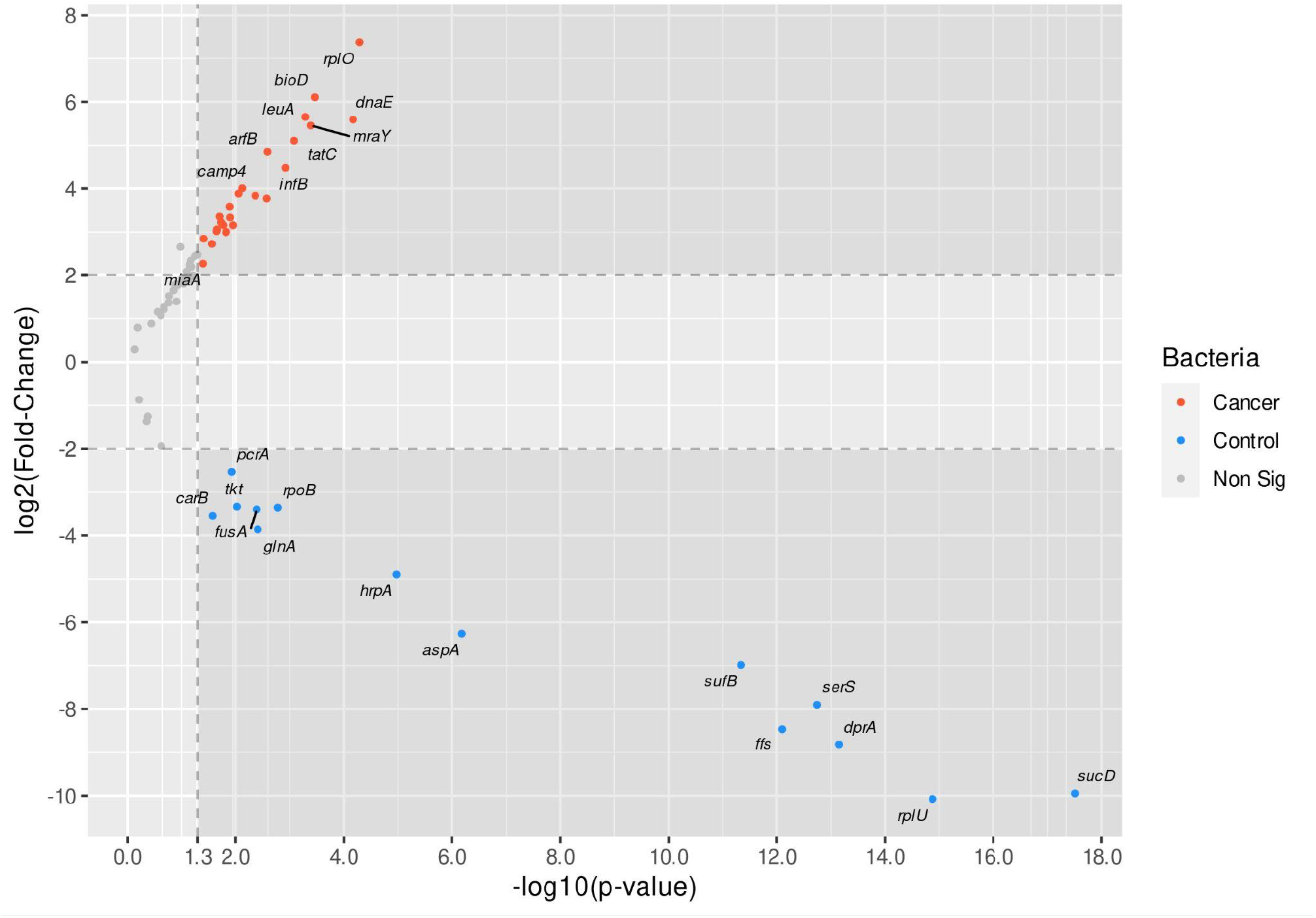
Differentially expressed bacterial genes

This reference uses the genomes of the most abundant microorganisms in the selected group. The number of genomes included is determined by the user and depends on the group chosen for analysis (all samples, case vs. control, or sample by sample). Once genes are quantified, several downstream analyses can be performed.

## 4. CONCLUSION

Regarding the methodology, RNA-Seq Total not only characterizes microbiome taxonomy similarly to 16S but also surpasses this approach by enabling functional analysis based on non-human RNA expression levels. Furthermore, the pipeline’s central taxonomic aligner, Kraken, demonstrates enhanced sensitivity through nucleotide-level alignment using complete genome sequences. Accurate identification of the microorganisms present in the environment enables downstream analyses of RNA-Seq data, including gene expression quantification, functional profiling, and differential analyses.

This conjecture arose during the analysis of a sample with known abundance, coupled with the evaluation of another study employing a RNA-Seq Total methodology. The present study advocates for the development and implementation of an analytical model capable of deriving insights beyond solely taxonomic analysis from the microbiome, thereby enriching the ongoing discourse within the scientific community on this emerging subject.

## APPENDIX I ARTICLES USED FOR THE LITERATURE REVIEW OF THE MOST PREVALENT TAXA BY SAMPLE TYPE

1. Tissue (Bacteria, Viruses, Fungi, and Archaea):
  a. *Gastric Cancer Microbiome*
  b. *The role of virome in the gastrointestinal tract and beyond*
  c. *Non-bacteria microbiome (virus, fungi, and archaea) in gastrointestinal cancer*
2. Feces (Bacteria, Viruses, Protozoa, and Archaea):
  a. *The microbiome and systemic sclerosis: A review of current evidence*
  b. *Assessment of Gastroenteric Viruses from Wastewater Directly Discharged into Uruguay River, Uruguay*
  c. *Fecal Influenza in Mammals: Selection of Novel Variants*
  d. *Molecular and biological characterization of influenza A viruses isolated from human fecal samples*
  e. *Survival of murine norovirus and hepatitis A virus in bottled drinking water, strawberries, and oysters*
  f. *The importance of fecal nucleic acid detection in patients with coronavirus disease (COVID-19): A systematic review and meta-analysis*
  g. *Commensal Intestinal Protozoa—Underestimated Members of the Gut Microbial Community*
3. Water (Bacteria, Viruses, Fungi, Protozoa, and Archaea):
  a. *A comparative analysis of drinking water employing metagenomics*
  b. *Microbiomes in drinking water treatment and distribution: A meta-analysis from source to tap*
  c. *Survival of murine norovirus and hepatitis A virus in bottled drinking water, strawberries, and oysters*
  d. *SARS-CoV-2 coronavirus in water and wastewater: A critical review about presence and concern*
4. Soil (Bacteria, Viruses, Fungi, Protozoa, and Archaea):
  a. *Influence of resistance breeding in common bean on rhizosphere microbiome composition and function*
  b. *Linking rhizosphere microbiome composition of wild and domesticated Phaseolus vulgaris to genotypic and root phenotypic traits*
  c. *Diversity and potential biogeochemical impacts of viruses in bulk and rhizosphere soils*
  d. *Dominant protozoan species in rhizosphere soil over growth of Beta vulgaris L. in Northeast China*
  e. *Isolation and identification of some fungi from rhizospheric soils of some wild plants at Iraq*
  f. *Bacteria, Fungi and Archaea Domains in Rhizospheric Soil and Their Effects in Enhancing Agricultural Productivity*
  g. *The rhizosphere zoo: An overview of plant-associated communities of microorganisms, including phages, bacteria, archaea, and fungi, and of some of their structuring factors*

## APPENDIX II TAXA BY SAMPLE TYPE

1. Tissue:
  1. Bacteria (GC): Helicobacter, Prevotella, Streptococcus, Halomonas, Pseudomonas, Sphingomonas, Lactobacillus, Shewanella, Acinetobacter, Corynebacterium, Bacillus, Neisseria, Leptotrichia, Veillonella, Bacteroides;
  2. Bacteria (HC): Prevotella, Agrobacterium, Sphingobium, Helicobacter, Porphyromonas, Halomonas, Neisseria, Zoogloea, Shewanella, Rhodobacter, Vogesella, Sphingomonas, Erythromicrobium.
  3. Virus: EBV, Cytomegalovirus, Myoviridae, Podoviridae, Siphoviridae, crAssphages (Bacteriófagos), Adenoviridae^*^, Herpesviridae^*^, Iridoviridae^*^, Marseilleviridae, Mimiviridae^*^, Papillomaviridae^*^, Polyomaviridae^*^, Poxviridae^*^, Anelloviridae^*^, Circoviridae, Parvoviridae^*^, Picobirnaviridae^*^, Reoviridae^*^, Caliciviridae^*^, Astroviridae^*^, Virgaviridae, Picornaviridae^*^, Retroviridae^*^, Togaviridae^*^, Alphaflexiviridae, Bromoviridae, Luteoviridae
  4. Fungi: Malasseziomycetes, Saccharomycetes, Aspergillus, Malassezia, Rhodotorula, Pseudogymnoascus, Kwoniella, Talaromyces, Debaryomyces, Moniliophthora, Pneumocystis, Nosemia, *Candida, Alternaria, Saitozyma, Thermomyces*
  5. Archaea: Methanobrevibacter smithii, Methanosarcina mazei, Methanomicrococcus blatticola, Halobacterium salinarum, Natrinema sp. J7-2, Methanobrevibacter oralis
2. Feces (Systemic Sclerosis):
  1. Bacteria DE (CA e HC): Bacteroides, Escherichia, Lachnospira, Prevotella, Roseburia, Veillonella, Bifidobacterium, Faecalibacterium, Fusobacterium, Erwinia, Clostridium, Lactobacillus, Streptococcus, Desulfovibrio
  2. Virus: Influenza, arbovírus, novovírus, rodovírus, coronavirus, Hepatitis A Virus (^*^)
  3. Protozoa: Entamoeba, Blastocystis, Dientamoeba fragilis
  4. Archaea: Methanobrevibacter smithii
3. Drinking Water:
  1. Bacteria: Methylibium petroleiphilum, Methyloversatilis, Polaromonas, Hydrogenophaga, Ramlibacter tataouinensis, Hylemonella gracilis, Propionibacterium acnes, Bdellovibrio bacteriovorus, Brevundimonas bacteroides, Acinetobacter lwoffii, Salmonella enterica, Alishewanella jeotgali, Alishewanella agri, Alishewanella aestuarii, Methylocystis, Afipia birgiae, Bradyrhizobium elkanii, Mycobacterium kansasii, Mycobacterium, Rhodococcus, Flavobacterium, Nitrospira, Bradyrhizobium, Acidovorax, Aquabacterium, Nitrosomonas, Massilia, Pseudomonas
  2. Vírus: Influenza, arbovírus, novovírus, rodovírus e corona
  3. Fungi: Chrysosporium queenslandicum, Malassezia restricta, Melampsora pinitorqua
  4. Archea: Candidatus Nitrosoarchaeum koreensis
  5. Protozoa: Acanthamoeba palestinensis, Acanthamoeba mauritaniensis
4. Soil:
  1. Bacteria: Pseudomonas, Bacillus, Cytophagaceae, Solibacteraceae, Acidobacteria, Actinobacteria, Chlamydiae, Chloroflexi, Cyanobacteria, Firmicutes, Planctomycetes, Proteobacteria, Verrucomicrobacteria, Dyadobacter, Streptomycetaceae, Nocardioidaceae, Rhizobiaceae
  2. Virus: Retroviridae, Nanoviridae, Phycodnaviridae, Mimiviridae, Microviridae, Podoviridae, Myoviridae, Circoviridae, Siphoviridae, Lymphocriptovirus, Aurivirus, Pradovirus, Papilomaviridae
  3. Fungi: Fusarium oxysporum, Aspergillus, Penicillium, Ascomycota, Alternaria, Zygomycota, Mucor, Rhizopus, Phytium
  4. Archaea: Sulfolobus, Haloarchaea, Methanosarcinaceae, Methanosaetaceae, Methanomicrobiaceae, Methanobacteriaceae
  5. Protozoa: Colpoda, Tachysoma, Oxytricha, Vorticella, Bodo

## Notes

### Competing Interest Statement

The authors have declared no competing interest.

